# Stripenn detects architectural stripes from chromatin conformation data using computer vision

**DOI:** 10.1101/2021.04.16.440239

**Authors:** Sora Yoon, Golnaz Vahedi

**Affiliations:** Department of Genetics, University of Pennsylvania Perelman School of Medicine, Philadelphia, PA 19104, USA; Institute for Immunology, University of Pennsylvania Perelman School of Medicine, Philadelphia, PA 19104, USA; Epigenetics Institute, University of Pennsylvania Perelman School of Medicine, Philadelphia, PA 19104, USA; Institute for Diabetes, Obesity and Metabolism, University of Pennsylvania Perelman School of Medicine, Philadelphia, PA 19104, USA; Abramson Family Cancer Research Institute, University of Pennsylvania Perelman School of Medicine, Philadelphia, PA 19104, USA

## Abstract

Architectural stripes tend to form at genomic regions harboring genes with salient roles in cell identity and function. Therefore, the accurate identification and quantification of these features is essential for the understanding of lineage-specific gene regulation. Here, we present Stripenn, an algorithm rooted in computer vision to systematically detect and quantitate architectural stripes from chromatin conformation measurements of various technologies. We demonstrate that Stripenn outperforms existing methods, highlight its biological applications in the context of B and T lymphocytes, and examine the role of sequence variation on architectural stripes by studying the conservation of these features in inbred strains of mice. In summary, Stripenn is a computational method which borrows concepts from widely used image processing techniques for demarcation and quantification of architectural stripes.

## Introduction

The eukaryotic genome is tightly organized inside the nucleus by forming a complex of DNA and histone proteins called the chromatin^1–3^. Chromosome conformation capture techniques, in particular Hi-C, suggest that the chromatin is folded into various length scales, forming a hierarchical structure^4,5^. Among these structures, topologically associating domains (TADs) are sub-megabase regions where stronger interactions are observed between loci inside each domain compared with loci in neighboring domains^6,7^. The spatial proximity of two genomic regions within TADs displaying high contact frequency in Hi-C maps is referred to as chromatin loops^8,9^. Loop formation, which is best described by the loop extrusion model, is mediated by sliding cohesin anchored by CTCF binding events in a convergent orientation^10,11^.

Although TADs and chromatin loops as distinguished features of Hi-C maps were described in pioneering studies, structures appearing as lines, flames or stripes attracted attention most recently^10,12–14^. Architectural stripes form through the loop extrusion process when a loop anchor interacts with the entire domain at high frequency^13^. Stripes have been reported as frequent features of diverse developmental programs^15^. It has been proposed that these unique structures, often associated with active enhancers and super-enhancers, can tether enhancers to cognate promoters, facilitating transcription and recombination^13^. Moreover, stripe anchors represent major hotspots for topoisomerase-mediated lesions, which may promote chromosomal translocations and cancer^13^. Since architectural stripes tend to form at genomic regions harboring genes with key roles in cell identity and function, it is essential to accurately detect these features from Hi-C or other chromatin conformation capture measurements^12,13,16^.

Numerous computational techniques have been developed to detect chromatin loops. Yet, the reliable identification of architectural stripes remains a challenge^17–20^. The first published stripe detection algorithm, referred to as Zebra, exploited the Poisson statistics and yielded thousands of stripes^13^. Despite the utility of Zebra in detecting strong regions with stripy features, the algorithm has three major limitations: (a) it has a high false positive rate and detects some chromatin loops as stripes, (b) it lacks a quantitative assessment of stripes and has been reported to rely on manual curation, and (c) the code for algorithm’s implementation is not publicly available.

Another computational approach, which is publicly available and referred to as domainClassifyR, was developed to comprehensively detect stripes by first marking TADs and then measuring their stripe and loop scores^12^. However, domainClassifyR assumes that stripes form exclusively at boundaries of genomic domains defined by TAD callers. It is evident from Hi-C maps of diverse cell types that although some architectural stripes form at TAD boundaries, stripy features are also found inside TADs. Hence, intra-TAD stripes remain undetected by the domainClassifyR computational approach. Another tool called CHESS was recently developed to perform quantitative comparisons of chromatin contact data between two conditions using the structural similarity index^21^. Although CHESS has been developed to report differential features (TADs, stripes or loops) between two conditions^21^, it cannot be used to delineate architectural stripes in a cell type of interest. Altogether, despite the availability of some computational techniques, there is a need to develop a quantitative strategy to accurately and efficiently detect architectural stripes and assess the strength of stripes across cell types and conditions.

Here, we report the development of a new stripe detection tool called Stripenn, which borrows concepts from computer vision and image processing. The backbone of our novel method relies on *Canny edge detection*^22^, one of the most popular edge detection algorithms in the mature field of computer vision. Stripenn can be applied to any type of chromatin conformation capture data such as Hi-C^23^, HiChIP^24^, and Micro-C^25^. Our method offers two scoring systems: *P*-value to filter low-quality stripes and *stripiness* to rank stripes based on the continuity of interaction signal. We found that Stripenn outperforms Zebra and domainClassifyR when these techniques are applied on an ultra-high coverage Hi-C data. Our systematic analysis of stripes from B and T lymphocytes using Stripenn revealed that the majority of stirpes were on the transcriptionally active compartment, architectural proteins mediating loop extrusion such as CTCF and proteins in the cohesin complex were highly bound at stripe anchors, and cohesin loader Nipbl and active enhancer modifications were highly enriched at stripe domains. To examine the effect of natural genetic variation on stripe formation, we compared stripes in T lymphocytes from two inbred mouse strains. Although hundreds of genes were differentially expressed between the two strains, most stripes were conserved and demonstrated comparable stripiness. Nonetheless, subtle changes in interaction within select stripes are associated with large differences in transcriptional outputs. Together, Stripenn, which is freely available on github^26^, is a specialized tool dedicated to stripe detection, enabling the systematic and quantitative analysis of stripes.

## Results

### Overview of the Stripenn algorithm

Stripenn systematically detects and quantitates architectural stripes from genome-wide chromatin conformation measurements. Our method applies principles from computer vision to detect genomic anchors that interact with entire domains at high frequency (Figure 1a). Since the algorithm treats contact maps as digital images, the performance of stripe detection depends on sequencing coverage and hence the resolution of contact matrix. The program’s output is a table containing coordinates and scores of the predicted stripes, which can be used for visualization purposes.

**Figure 1.**
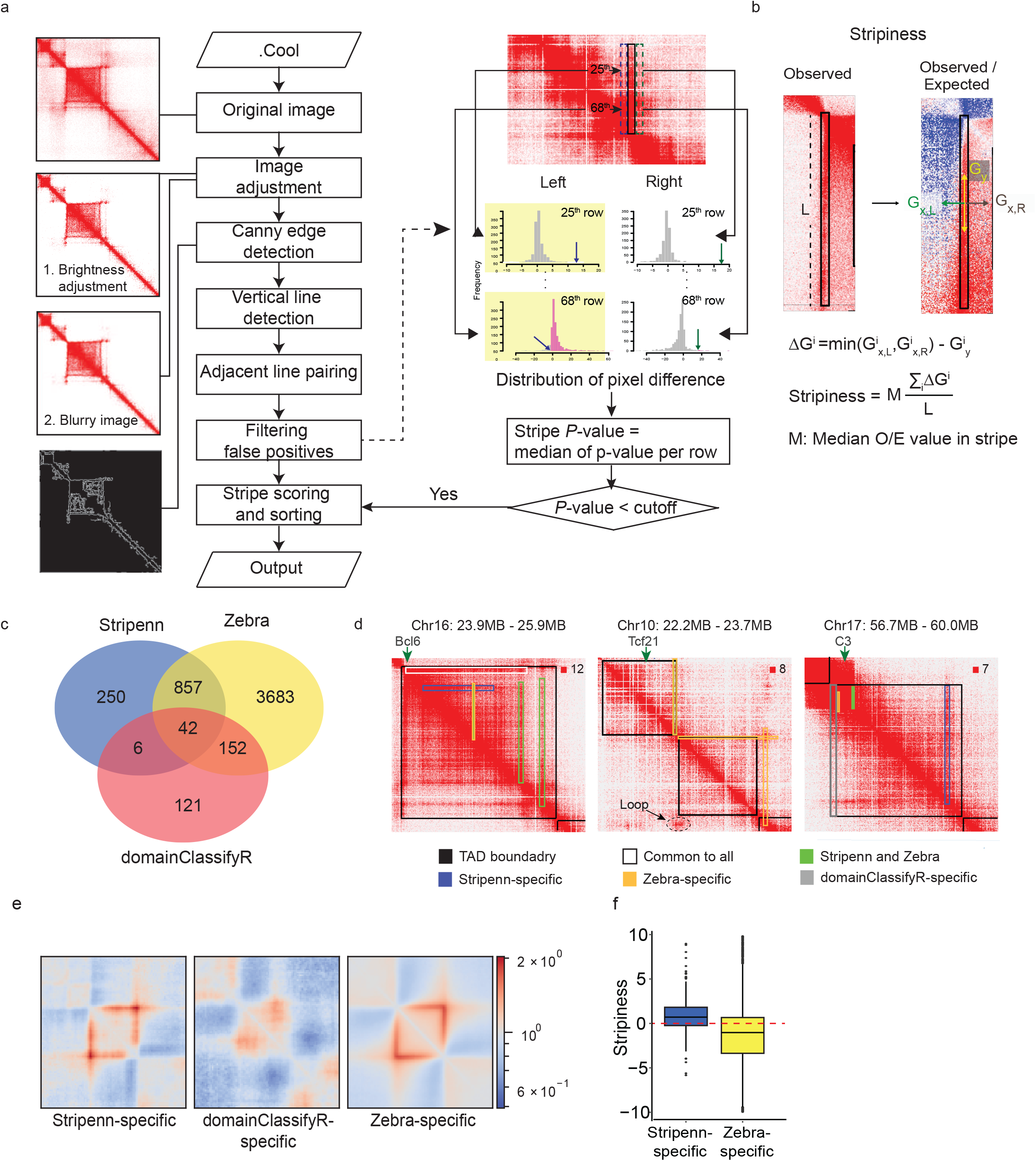
Stripenn overview and comparison with Zebra and domainClassifyR. (a) Stripenn searches for candidate stripes using image-processing techniques (left) and filter them based on median *P*-value calculation which estimates stripes contrast (right). *P*-value is estimated for every row in a stripe and the median is used as a median *P*-value. For each row, *P*-value is estimated based on the pixel difference within stripe (black border) and its left- or right-adjacent background (blue and green border, respectively). Among two p-values estimated from left and right backgrounds, the largest *P*-value is chosen (yellow box). Since the expected contact frequency decreases as genomic distance of two DNA regions increases, different null distribution is used for each row. A mock example of 25^th^ and 68^th^ rows is provided. (b) Filtered stripes are further ranked using stripiness measured based on observed/expected (O/E) contact frequency matrix. 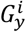: gradient in y-direction of i^th^ row. 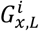 and 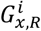: gradient in x-direction of i^th^ row (left and right direction, respectively) : Δ*G*^*i*^ refers to the difference of gradient in y-direction and x-direction. *L* refers to the length of stripe. (c) Venn Diagram represents the number of stripes detected by Stripenn, Zebra and domainClassifyR. (d) Representative stripes detected by Stripenn, Zebra and/or domainClassifyR are demarcated on Hi-C contact matrices. Black borders represent TAD boundary. (e) Pileup plots demonstrate quality of stripes exclusively detected by Stripenn, domainClassifyR and Zebra. (f) The distribution of stripiness of Stripenn-specific and Zebra-specific stripes.

The first step of Stripenn is to convert each chromosome’s contact matrix, which is provided in the cooler file format^27^, to a digital image. Next, contrast adjustment and noise reduction are applied to a sliding window of 500 pixels along a diagonal line of the image (Figure 1a). To enhance the contrast between signal and background, brightness is adjusted for multiple levels. To reduce noise, Gaussian blur effect is applied to each brightened image. Then, Canny edge detection algorithm^22^, which is the most popular edge detection technique to date, is applied to these processed images. The Canny edge detection algorithm, which is the backbone of Stripenn, relies on computing the gradient intensity of an image and applying various gradient magnitude thresholding to minimize spurious response to edge detection (see Methods). An edge in any digital image can point to a variety of directions; hence, the Canny algorithm uses filters to detect horizontal, vertical and diagonal edges. Considering that contact matrices are symmetrical about the diagonal line, Stripenn only reports vertical edges predicted by the edge detection algorithm. Since a stripe is defined by a genomic anchor positioned on the diagonal line, predicted vertical edges longer than 10 pixels are connected to the diagonal line. Canny edge detection predicts two edges for each stripe and adjacent lines within a pre-defined distance are paired, forming a single vertical stripe. Together, Stripenn adapts principles from digital image processing and edge detection techniques to demarcate architectural stripes.

### Two scoring systems: *P*-value and stripiness

To quantitate an architectural stripe, we devised two scoring systems: median *P*-value of pixel contrast and *stripiness* (Figure 1a-b). The median *P*-value is the median of *P*-values of rows in a stripe and is used to evaluate the contrast between a predicted stripe and its neighbors (Figures 1a and S1a). Each *P*-value represents the significance of the pixel differences between a given row and its left or right neighboring pixels. To reduce the noise effect, we applied data smoothing by calculating mean intensity of interactions in a window of 50kb height around a given stripe row. Moreover, the mean intensity of left (*L*) and right (*R*) neighboring pixels are calculated from 50 kb × 50 kb windows adjacent to a given row. To estimate the *P*-value of the difference between the mean intensity of neighboring windows from the center (*C* − *L* and *C* − *R*), a null distribution of the pixel difference is created for 1000 random genomic regions. Since the expected contact frequency decreases for long-range interactions, null distributions are separately created from one to 400 pixels away from the diagonal line. To avoid overestimating the median *P*-value, the less significant value is chosen among left and right *P*-values. Hence, the median *P*-value is a metric of contrast and is used to filter low quality stripes.

Although the *P*-value calculation examines the contrast between a predicted stripe and its neighboring pixels, this metric is not capable of assessing the continuity of interactions within such structures. To penalize discontinuous stripes which may represent loops, we devised another metric referred to as *stripiness* (Figure 1b). Stripiness relies on subtraction of the gradients in vertical and horizontal directions and considers the contrast between a stripe and its neighbors, pixel continuity within a stripe, as well as a stripe’s median pixel value (See Methods). Together, our new algorithm exploits *P*-value and stripiness as scoring criteria to quantitate intensity, continuity, and contrast of predicted architectural stripes.

### Benchmarking of Stripenn, Zebra, and domainClassifyR

The principles of Canny edge detection algorithm engineered in Stripenn can be easily applied to chromatin conformation measurements of different technologies including Hi-C^23^, HiChIP^24^, and Micro-C^25^ (Figure S1b). A larger number of stripes was predicted from Micro-C compared with Hi-C and HiChIP, suggesting that Micro-C can enable us to investigate more detailed chromatin structures that may be functionally relevant to gene regulation. The larger number of predicted stripes from this assay is consistent with the notion that Micro-C can overcome the current resolution gap of Hi-C at the fine scale^25^. To systematically compare Stripenn with existing methods including Zebra^13^ and domainClassifyR^12^, we applied these algorithms to the ultra-deep coverage Hi-C data in activated B cells^13^. This comparison revealed that domainClassifyR, Stripenn, and Zebra could detect 321, 1162, and 4781 stripes, respectively, from the same dataset (Figure 1c). Representative examples of predicted stripes portrayed on Hi-C contact maps were indicators of sensitivity and specificity of these techniques (Figure 1d). The most prototype stripes at TAD boundaries, such as the one harboring *Bcl6*, were detected by the three methods (Figure 1d, white stripe in left panel). However, intra-TAD stripes, which consist of more than 70% of all stripes (Figure S1c) and mostly have positive stripiness values (Figure S1d) could not be detected by domainClassifyR. Representative examples of top-ranked domainClassifyR-specific stripes did not demonstrate strong continuous interactions at these predicted loci (Figure 1d, grey stripe in right panel). Although Zebra appeared to be a sensitive technique predicting many stripes, the algorithm sacrificed specificity since loops or corner dots with weak stripy features were occasionally predicted as stripes (Figure 1d, yellow stripe middle panel). To complement the visual inspection of representative stripes, we next systematically compared the quality of stripes predicted by the three tools. First, we calculated the stripiness of Stripenn and Zebra predictions. This comparison revealed that while most common stripes between Zebra and Stripenn in addition to Stripenn-specific stripes showed positive stripiness (Figure S1e), the average stripiness of Zebra-specific stripes was negative, indicating the low quality of these predictions (Figure 1f). Of note, stripiness of Stripenn-specific stripes was slightly lower than those structures predicted by both Zebra and Stripenn (Figure S1e). Stripiness of stripes predicted by domainClassifyR could not be evaluated because this technique reports TAD coordinates and does not provide information on the exact genomic positions of stripes. Moreover, the pileup plot analysis showed that most domainClassifyR-specific predictions did not form stripe features compared to Stripenn- and Zebra-specific predictions (Figure 1e). In contrast to Zebra-specific stripes, which demonstrated poor contrast between predicted stripes and their surroundings in pileup plots, the Stripenn-specific stripes had a significantly higher contrast represented by the lower level of contacts around genomic regions adjacent to predicted genomic coordinates. Together, Stripenn outperforms two existing stripe callers and can quantitate the intensity, continuity, and contrast of predicted architectural stripes.

### Architectural stripes are favored on transcriptionally active regions

The relevance of architectural stripes to gene expression has been examined in studies which utilized Zebra^13^ or domainClassifyR^12^ as stripe callers. To evaluate Stripenn’s performance across different technologies, we first investigated Stripenn’s predictions on Hi-C measurements from activated B cells. Using median *P*-value < 0.05, stripiness > 0, and 5kb-resolution, we found 537 stripes in activated B cells (390 5’- and 147 3’-stripes). We examined the compartmentalization of stripes and found that stripes were frequently (~82%) formed on the transcriptionally active A compartment, which is consistent with the previous report using Micro-C measurements relying on stripy features at TAD boundaries^25^. Only 1.1% of the stripes were detected in the B compartment and around 16% of stripes spanned A and B compartments (Figure 2a). This selective enrichment of stripes on A compartment is in contrast with compartmentalization of TADs where only half of TADs (44.2%) are positioned in the A compartment, emphasizing the association of stripes with the active chromatin state. These findings are consistent with the observation that stripes frequently accommodate super-enhancers^12,13^. We further focused on TADs harboring stripes, referred to as “stripy” TADs, and compared them with those without any stripes, referred to as “non-stripy” TADs. We examined genomic length and the enrichment of architectural proteins including CTCF, cohesin subunits Smc3, Rad21, and cohesin loader Nipbl in two TAD classes. The stripy TADs were defined as TADs containing any stripe with median *P*-value < 0.05 and stripiness > 0, and the non-stripy TADs did not include any stripes predicted by Stripenn and only TADs on the A compartment were considered in this comparison. Together, 155 stripy and 286 non-stripy TADs were considered for further analysis. Interestingly, we found that stripy TADs were significantly longer in genomic length compared with non-stripy TADs (Figure S2a). TADs in the A compartment are on average smaller in genomic length compared with those on the B compartment (Figure S2c-d) and some stripes span both A and B compartments (Figure 2a). Hence, we compared the genomic length of stripy and non-stripy TADs in the A compartment and found that the disproportionate difference in genomic length was also present when stripy TADs exclusively in the A compartment were considered for this comparison (Figure S2b). Together, the link between genomic length and stripe formation on active compartment may imply the accommodation of numerous regulatory elements in stripy TADs. We next compared the median intensity profiles of architectural proteins on stripy and non-stripy TADs and found that architectural proteins such as CTCF, Smc3, and, Rad21 had higher intensity of binding on stripy TADs compared with non-stripy TADs (Figure S2 e-f). Moreover, consistent with previous studies^12,13^, CTCF and cohesin subunits were highly enriched at stripe anchors (Figures 2a and S2g). These data suggest that stripes favor the transcriptionally active A compartment and the patterns of architectural proteins binding at stripy TADs may implicate their roles in stripe formations.

**Figure 2.**
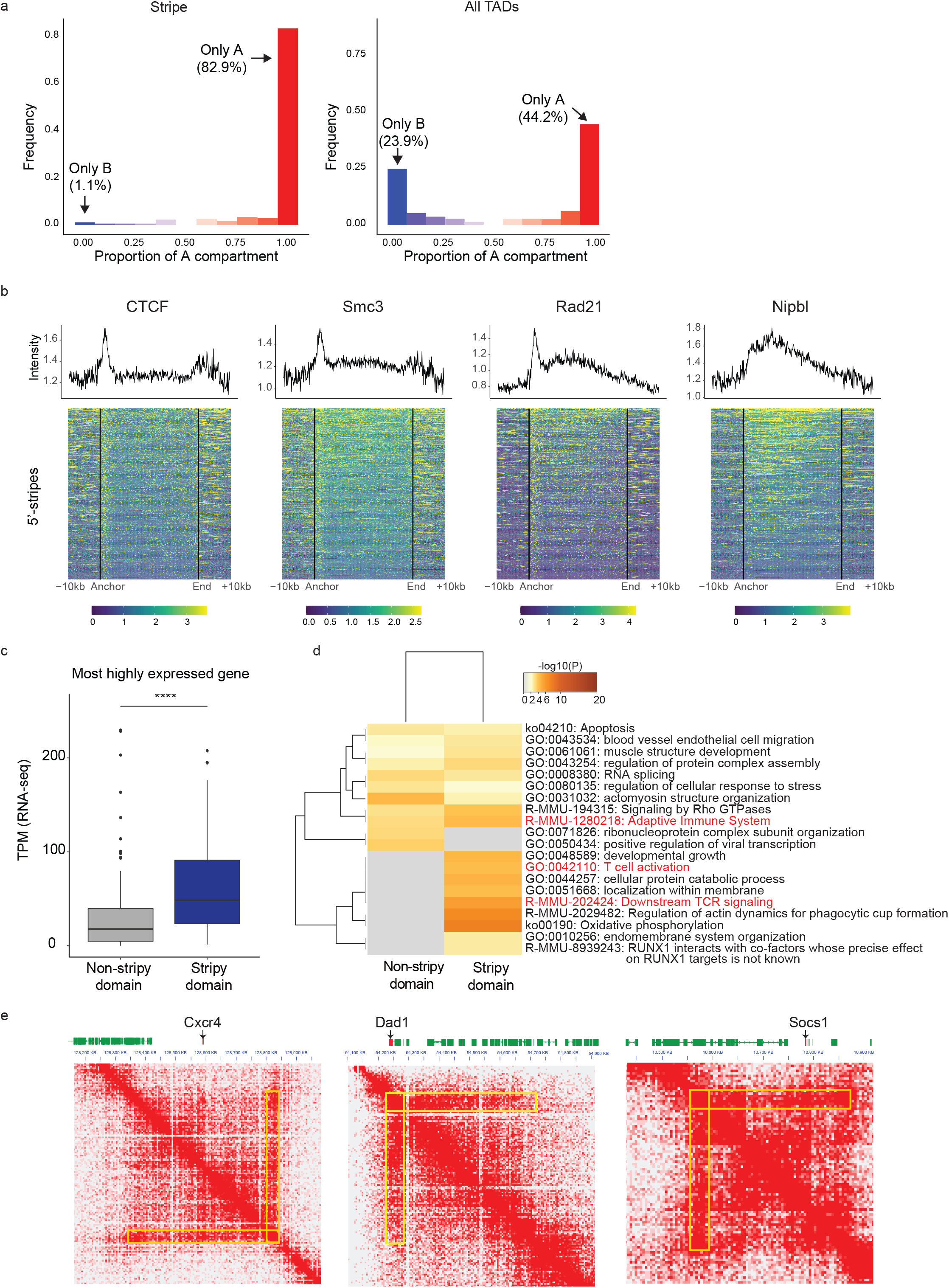
Stripes are located on transcriptionally active regions and associated with cell-type specific gene expression. (a) Genomic fraction of stripe domain overlapping with A compartment. Majority of stripes (82.9%) are located on the A compartment (left) unlike TADs which are distributed evenly between two compartments (right). (b) Distribution of CTCF, Smc3, Rad21 and Nipbl on rescaled 5’-stripes. The profiles represent the median intensity of each proteins. Black lines in each heatmap are boundaries of stripes. (c) Comparison of the most highly expressed gene expression levels between stripy (blue) and non-stripy (gray) TADs. (d) The gene ontology analysis of the most highly expressed genes in stripy and non-stripy TADs using Metascape. Immune response-related terms are marked red. (e) Examples of Thymocyte-specific genes (*Cxcr4, Dad1* and *Socs1*) included in stripes from DP thymocyte.

### Stripy TADs are more accessible and possess more active enhancers than non-stripy TADs

To further assess the quality of architectural stripes detected by chromosome conformation techniques other than Hi-C (Figure S1b), we applied Stripenn to HiChIP measurements. HiChIP is a ligation-proximity reaction assay, which detects interacting DNA fragments bound by a protein of interest^24^. We applied Stripenn to our recently generated Smc1 HiChIP in double positive (DP) thymocytes in C57BL/6J mice^28^ and detected 431 stripes (314 5’- and 117 3’-stripes) (median *P*-value < 0.05 and stripiness >0). Consistent with findings based on Hi-C maps of B cells, we confirmed that most stripes formed on the A compartment of DP thymocytes (Figure S3a). Moreover, we found that deposition of active enhancer mark, H3K27ac, was skewed towards stripe anchors (Figure S3b). Furthermore, the active enhancer marks were significantly more enriched at stripy TADs compared to non-stripy TADs (Figure S3c). In addition, stripy TADs were on average more accessible than non-stripy TADs (Figure S3d). Together, these data further corroborate that architectural stripes favor highly accessible and active chromatin states.

### Stripy TADs are formed at highly expressed genes associated with T cells

To assess the association between stripe formation and transcriptional outputs, we compared the expression levels of genes located on 156 stripy TADs and 144 non-stripy TADs within the A compartment of thymocytes. On average, we did not find any significant difference between the expression levels of genes located on stripy vs non-stripy TADs (Figure S3e). Nonetheless, we found that the most highly expressed genes encompassing stripy TADs were significantly more expressed than the most highly expressed genes encompassing non-stripy TADs (Figure 2c). This finding suggests that not all but distinct genes in stripy TADs benefit from the continuous interaction of a stripe anchor with regulatory elements located on a genomic domain. To further delineate the identity of most highly expressed genes, we performed gene ontology analysis using Metascape^29^. Interestingly, genes located on stripy TADs were more significantly enriched in the immune response-related terms such as ‘adaptive immune system’, ‘T cell activation’ and ‘downstream TCR signaling’, compared with genes on non-stripy TADs (Figure 2d). Representative examples demonstrate the interactions of stripe anchors and T cell associated genes such as *Cxcr4*, *Dad1*, and *Socs1* (Figure 2e). The gene encoding the chemokine receptor, CXCR4, which is recruited into immune synapse during T-cell activation^30^, is in the middle of a stripy TAD and can act as a boundary of two nested TADs. Another representative architectural stripe is located on an anti-apoptosis gene *Dad1,* which can enhance T-cell proliferation^31^, and is located downstream of the T-cell receptor alpha chain gene (TCR*α*)^32,33^. Similar to *Cxcr4*, *Socs1* required to suppress the cytokine signaling and regulate T cell proliferation, activation, and function^34^, resides in the middle of a stripy TAD and is preferentially located near the boundary of two nested TADs. The selective enrichment of T cell associated genes within architectural stripes of T cells further confirms the potential cell type-specific regulatory role of stripes.

### Stripes are strongly conserved between two inbred mouse strains

We next aimed to investigate whether genetic perturbations can alter stripe formation and be linked to transcriptional regulation. Hence, we relied on millions of natural genetic variations between two inbred mouse strains C57BL/6J and nonobese diabetes (NOD) mice. Using Smc1 HiChIP measurements in DP thymocytes of these mice, Stripenn was able to detect 953 and 1151 stripes in C57BL/6J and NOD mice, respectively^28,35^. To compare the degree of interactions within stripes between two strains, stripiness and median *P*-values of stripes merged across two strains were calculated. Despite more than 5 million single-nucleotide polymorphisms and 400 thousand insertions and deletions, the stripiness of predictions in DP thymocytes of two mouse strains was significantly correlated (Figures 3a and S4a). The visual inspection of loci encompassing major T cell associated genes such as *Bcl6* and *Ets1* demonstrated a large-scale conservation of architectural stripes between two strains (Figure S4b-c).

**Figure 3.**
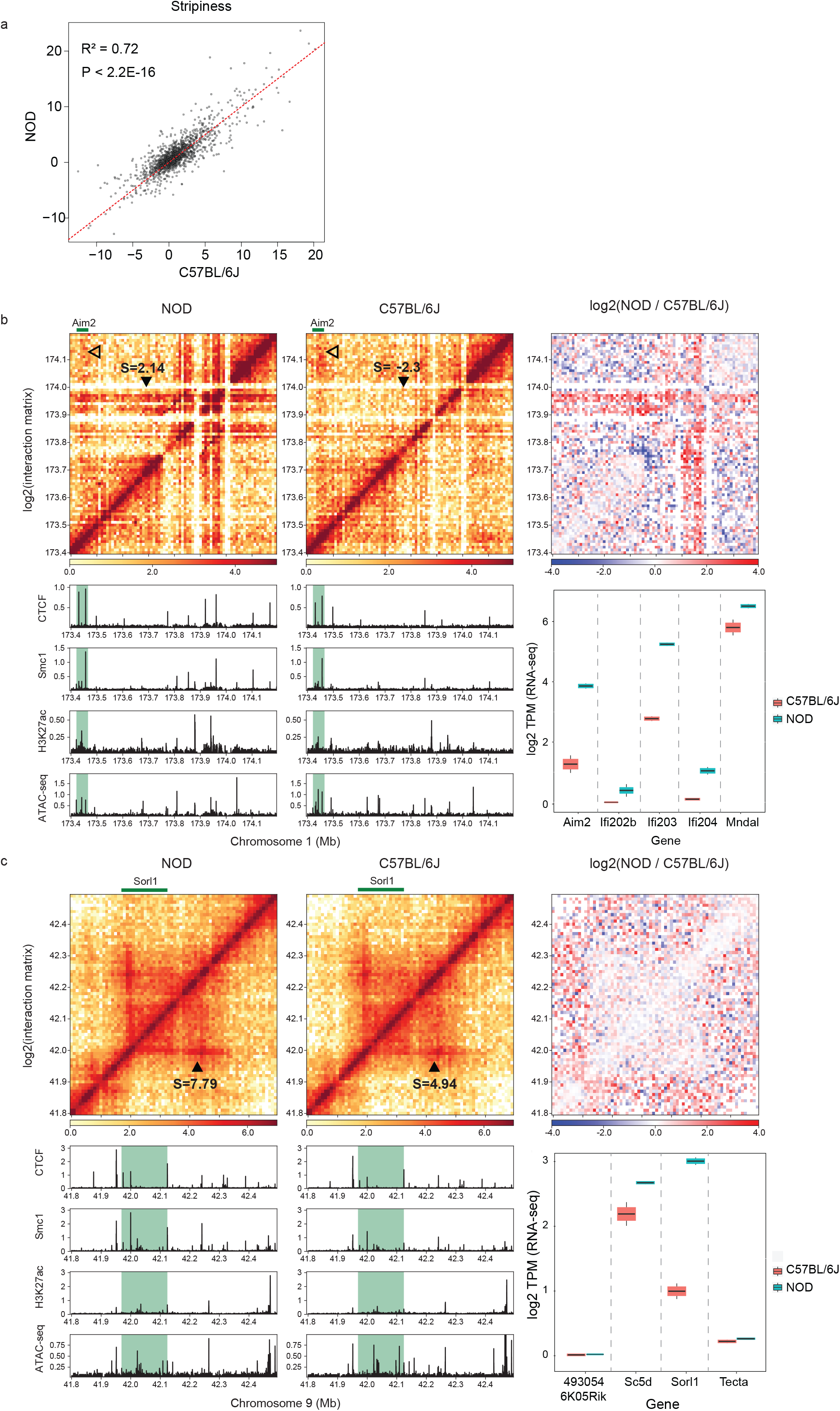
Stripes are mostly preserved between T-cells of two inbred mouse strains. (a) Stripes were extracted from Smc1-HiChIP data of DP thymocyte of control (C57BL/6) and prediabetic (NOD) mice. Stripenn predictions in two strains were merged and then stripiness of all stripes was recalculated based on C57BL/6J and NOD HiChIP data. (b) A representative example demonstrates a differential stripe between NOD and C57BL/6J strains. Stripiness (*S*) of differential stripe is represented using triangles. Empty triangle indicates a conserved stripe that have Aim2 as its anchor. Contact frequency heatmap, ChIP-seq intensity of CTCF, Smc1, H3K27ac and ATAC-seq signal on Aim2 stripe region is shown for NOD and C57BL/6J. The heatmap in the third column demonstrates contact frequency ratio between two strains. The boxplot represents the expression level (TPM) of genes located in the stripe domain in NOD (blue) and C57BL/6J (pink). (c) An example of conserved stripe across two strains including a differentially expressed gene (Sorl1). Stripiness (*S*) of two stripes are represented in the heatmap.

Despite this global similarity of stripes between two strains, the stripiness metric demonstrated exceptional stripes with enhanced interactions in a strain-specific manner. One example is a stripe specific to NOD strain harboring killer cell lectin-like receptor (*Klr*) gene family located on chromosome 6 (Figure S4d). These genes, which are typically expressed in natural killer cells, identify and enable the destruction of virus-infected cells^36^. Our own group recently showed that this cluster of genes forms a set of long-range genomic interactions among enhancers and promoters, also referred to as hyperconnected 3D cliques^28^. These genes were lowly expressed in T cells of C57BL/6J but showed consistently higher expression in NOD mice (Figure S4e). In addition, the intensity of CTCF, Smc1 and H3K27ac as well as DNA accessibility were much higher in the NOD strain (data not shown). Another NOD-specific stripe contained genes such as *Aim2*, *Ifi202b*, *Ifi203*, *Ifi204,* and *Mndal* on chromosome 1. Protein products of *Aim2* are involved in inflammasomes and act as sensors for pathogen or abnormal DNA in the cytoplasm^37^. Stripenn predicted *Aim2* as a stripe anchor in thymocytes of both strains but with insignificant median *P*-value and stripiness (Figure 3b). However, a NOD-restricted stripe crossing *Aim2* was predicted by Stripenn and the visual inspection of this locus confirms significantly enhanced interactions in NOD compared with C57BL/6 strain. Moreover, CTCF, Smc1, and H3K27ac modification were significantly more abundant at the anchor of the NOD-specific stripe only in this strain. Consistent with enhanced chromatin interactions at this locus, all genes in this region showed higher expression levels in NOD compared with C57BL/6J (Figure 3b).

### Subtle changes in local interactions are linked to differential gene expression

Since majority of stripes were conserved between two strains, we next focused on stripes encompassing differentially expressed genes. The differential gene expression analysis yielded 359 and 336 genes upregulated in NOD and C57BL/6J, respectively (adjusted *P* value < 0.025, log2 fold change > 0.5). Nonetheless, we were not able to detect a significant difference in interactions of stripy regions between the two strains accommodating differentially expressed genes (Figure S4f). A representative example at the *Sorl1* locus demonstrates an exception to this trend. *Sorl1* is highly expressed in thymocytes of NOD compared with C57BL/6J mice (adjust *P* value=2.09e-17). Although the architectural stripe formed at this locus was detected by Stripenn in both strains, the stripiness of this stripe was ~2 times higher in NOD compared with C57BL/6J (Figure 3c; highlighted region). The stronger enrichment of Smc1 might have increased enhancer and promoter interactions in this region, leading to higher expression levels of *Sorl1* in NOD mice^38^. This example further suggests that subtle changes in local chromatin interactions, which are detectable by Stripenn, may be linked to changes in gene expression.

## Discussion

Since its invention in 2009^23^, Hi-C and its variations suffered from the unreasonably high cost of sequencing required to map high-resolution features such as enhancer-promoter interactions. With the cost of sequencing dropping significantly due to sequencing platforms such as NovaSeq and the emergence of single-nucleosome resolution techniques such as Micro-C^25^, the 4D nucleome community has witnessed the emergence of fine structures such as architectural stripes, which are not detected in low-resolution 3D genome measurements. Despite the pioneering work which suggested that cohesin continually extrudes loops of chromatin *in vivo* to form architectural stripes relying on ATP to fuel loop extrusion^13^, detailed molecular processes through which some regions form stripes, but not stable loops remain unclear. A major technical barrier to study these features and compare them across various developmental programs is the lack of a computational method to accurately demarcate and quantitate architectural stripes. Here, reasoning that stripes resemble edges of a digital image, we developed a novel technique with high accuracy and sensitivity relying on the most popular and widely-used edge detection algorithm in computer vision. The comparison of this novel technique with previously developed stripe callers revealed that relying on edge detection algorithms enables the identification of accurate architectural features. Consistent with previously published studies, we demonstrated that architectural stripes are enriched at transcriptionally active and accessible genomic regions. Moreover, we mapped these features in inbred strains of mice and found the large-scale conservation of architectural stripes despite millions of single nucleotide variation, suggesting the resilience of architectural stripes to sequence variation.

An intriguing feature of architectural stripes relates to their cell-type specificity and the preferential formation at lineage-determining genes. A critical yet unanswered question relates to mechanistically distinguishing stripes from chromatin loops. It remains unclear how architectural stripes selectively form in a particular cell type but can represent as corner dots or chromatin loops in other cell types. It remains unlikely that clustering of CTCF recognition sites and their orientation can play a role considering the identical underlying DNA sequence in different cell types. Whether cell-type specific CTCF binding events dictated by cell-type specific accessible chromatin regions at stripe anchors or additional epigenetic modifications such as DNA methylation can earmark the formation of stripes in select cell types remains to be shown. Exploiting machine-learning strategies and using large-scale detection of architectural stripes by Stripenn across different cell types collected by the 4D nucleome community can pave the way to better understand the grammar of stripe formation.

## Methods

### Stripenn

Stripenn detects architectural stripes from chromatin conformation capture data using image processing principles and score them with median *P*-value and stripiness. Stripenn can be installed via pip (or pip3) and more information of installation and usage are described in the GitHub page (https://github.com/vahedilab/stripenn).

#### Input and output

The input of Stripenn is in a cooler file format (.cool or .mcool) that contains genomic matrix such as Hi-C contact frequency matrix^27^. The output is a table of stripe coordinates, sizes, and measures (median *P*-value, stripiness and average/sum of pixel values within stripes). Stripenn runs through three steps to generate the output: (1) to search candidate stripes, (2) to measure median *P*-value, and (3) to measure stripiness.

#### To search for candidate stripes

For each chromosome, stripes are searched within a series of windows of 400 pixels that is moving along a diagonal line of contact frequency heatmap using the step size of 200 pixels. In each window, referred to as submatrix hereinafter, rows and columns composed of only zeros are removed and the corresponding genomic regions are deleted. Stripenn follows eight steps to find candidate stripes in each submatrix:

1. *Convert matrix to image:* To detect stripes, a submatrix is converted to an image as shown in Figure 1a. In this process, pixel values in a submatrix should be truncated to an appropriate value, which we refer to as *maximum pixel value*, to visualize the 3D chromatin structure properly. This step is important since extremely low or high maximum pixel value makes the image covered with only red or white pixels, which does not give any information. Because the stripe detection is sensitive to the maximum pixel value, Stripenn can search stripes for multiple maximum pixel values. These are determined by the percentiles of contact frequencies that the users provide as input. For example, the option ‘−m 0.93,0.95,0.97’ in the command line enables to set the maximum pixel values as top 7%, 5% and 3% of the positive contact frequency values of each chromosome. These percentiles should be set differently for different data based on the sequencing depth of the data. Stripenn’s ‘*seeimage*’ function helps the users decide the percentile values by visualizing a contact frequency matrix of given coordinates for given percentile. Once maximum pixel value (*M*) is determined, each pixel value (*P*) in submatrix is then converted to RGB codes such as

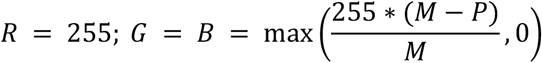 These RGB codes are then merged using *merge* function in opencv-python package^39^ to create image.
2. *Brightness adjustment*: Next, we adjust the brightness to increase the contrast between signal and noise of the image. Because the optimal brightness that best reveals the 3D structure in the image is not known, we adjust the brightness for multiple levels and find stripes for each level. To do this, MATLAB’s *imadjust* function^40^ is implemented in python and used with changing *high_in* parameters from 0.5 to 1.0 increasing by 0.1.
3. *Blur effect*: Next, blur effect was applied to reduce the noise of the image. Here, *filter2D* function of opencv-python package was used and 3 × 3 two-dimensional matrix filled with 1/9 was used as kernel.
4. *Canny edge detection*: The blurry image is then converted to gray scale using *cvtColor* function in opencv-python package to apply Canny edge detection. It is the most widely used edge detection method because it searches optimal edges that satisfies three criteria of an optimal edge: (1) edge with low error rate, (2) optimal localization and (3) only one strong signal is detected. The edge image ***E*** is extracted using *canny* function (sigma=2.5) in Scikit-image package^41^.
5. *Vertical line detection*: Because Hi-C data is symmetric about the diagonal line, horizontal stripes are automatically detected if corresponding vertical stripes are found. Thus, we search for only vertical stripes here. To do this, pixels whose orientations are not between 60° and 120° are removed. The orientation *θ* of each pixel ***E***(*i*, *j*) is calculated using Sobel operator as follows^42^.

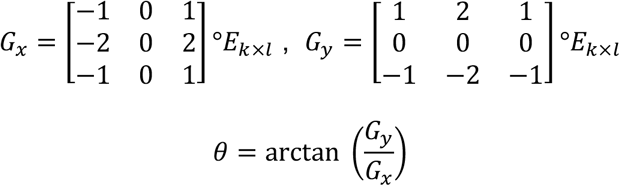 Where *G*_*x*_ and *G*_*y*_ are gradient in x- and y-direction, and *E*_*k×l*_ is 3 × 3 submatrix of edge matrix ***E***(*i* − 1 ≤ *k* ≤ *i* + 1, *j* − 1 ≤ *l* ≤ *j* + 1). Here, *convolve2d* function in SciPy package^43^ and *arctan2* function in NumPy package^44^ are used. Because the orientations of edges in *i*^*th*^ column are reflected in (*i* + 1)^*th*^ column of the orientation matrix, we shift the vertical line to 1 pixel left. Next, continuous pixels longer than 10 pixels are detected. Since the edges are usually not straight in real, we allow each pixel in a straight line to shift left or right by 1 pixel. These continuous lines are then extended to the diagonal line.
6. *Line refinement*: After step 5, several lines can be combined together since we allow 1 pixel shift. To find the representative line among them, we first converted pixel one (signal) in each line to zero (background) if it is originally zero in the edge image ***E***. Then, we find the representative column as the average column indexes weighted by their line lengths.
7. *Adjacent line pairing*: Next, we paired two vertical lines if their distance is less than *N* pixels (default: 8 pixels). This maximum stripe width should change according to the data resolution. For example, we recommend using 8 pixels and 16 pixels as maximum widths for 10kb and 5kb, respectively.
8. *Merging stripes*: Searching stripes from overlapping submatrices with multiple brightness parameter results in several duplicated stripes. At the last step of candidate stripe search, these are merged into one as the longest stripes.

#### Median P-value

The median *P*-value is devised to filter candidate stripes based on the pixel contrast between stripes and their backgrounds. Simply, the median *P*-value is a median of a series of *P*-values evaluated from each stripe row. The *P*-value of each stripe row is estimated as described in Figure S1a. First, mean contact frequency (*C*) of a given row is calculated. Here, we applied data smoothing to reduce the noise by deriving the mean contact frequency from 2D matrix of *N* × width pixels. Here, *N* is the number of pixels corresponding to 50kb and it depends on the resolution of data. Next, we calculate the mean contact frequency of both left (*L*) and right (*R*) neighborhoods of the testing row. The mean frequency is calculated from 2D matrix of *N* × *N* pixels right next to the testing row. Since the *P*-value estimates the significance of *C* − *L* and *C* − *R*, corresponding null distribution was constructed by randomly selecting 1000 data points. Considering that the bin distance affects the contact frequency, we selected data points of identical bin distance as testing row pixel (first pixel for 5’-stripe and last pixel for 3’-stripe). The null distributions of left and right neighborhoods are constructed separately so that the significance of *C* − *L* and *C* − *R* are accurately measured. To avoid overestimating the median *P*-value, we choose the less significant value. Stripenn output reports a list of stripes of which *P*-value is less than 0.1.

#### Stripiness

Median *P*-value measures the significance of contrast between stripe and the background; however, this metric does not reflect the contact frequency and continuity of intensity within stripe. To make up these shortcomings of median *P*-value, stripiness is devised. Stripiness (S) is estimated on observed/expected contact frequency matrix and it is defined as

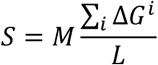

where *L* is the number of stripe rows, *i* is the index of stripe row (*i* = 1~*L*), *M* is the median of mean frequencies of stripe rows and Δ*G*^*i*^ is gradient change at *i*^th^ stripe row. The gradient change Δ*G*^*i*^ represents the difference between gradient in x-direction (across stripe and background) and y-direction (within stripe). Since larger gradient in x-direction and smaller gradient in y-direction are features of strong stripes, we define Δ*G*^*i*^ as

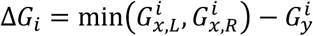

where 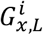 and 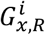 are gradient in left and right directions in *i*^*th*^ row, respectively. These gradient values are calculated using modified Sobel gradient as follows.

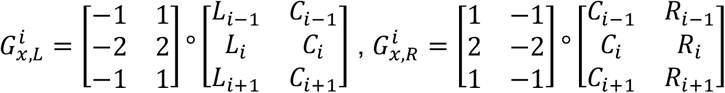

where *C*_*i*_, *L*_*i*_ and *R*_*i*_ are the mean of contact frequencies in *i*^th^ stripe row and adjacent left/right background rows (width=50kb), respectively. The gradient in y-direction 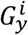 is calculated as.

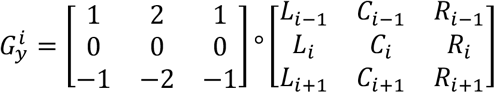

### Benchmark analysis

For competitive benchmarking, Hi-C data of 72 hours-activated B cell (GSE82144) was used. Zebra stripe calls were directly obtained from the author of the original paper^13^. These calls were processed so that adjacent calls are merged, and each call is connected to the diagonal line of the contact frequency matrix. To run domainClassifyR^12^, its R package (domainClassifyR) as well as misha^45^ and shaman^46^ packages were used. TAD coordinates, which are essential for this program, were assessed using two perl scripts ‘*matrix2unsulation.pl*’ and ‘*insulation2tads.pl*’ from cworld-dekker Github page. All parameters were set as defaults. Then calls with forward or reverse z-score > 5 were used in the analysis to have enough calls for comparison. Stripenn stripes were called from unnormalized, 10kb-resolution data to make it comparable to Zebra calls.

1. *Stripe overlap analysis:* Next, overlap among three stripe calls was counted. We regarded two stripes overlapping if they shared at least one genomic bin. If two or more stripes from a method were overlapping with a stripe from the other method, we counted it as one overlap.
2. Pileup plot analysis: Pileup plot of stripes that are unique to each method was generated using coolpup.py program^47^ where ‘--local --rescale’ options were used.

### Example chromatin conformation capture data

The example stripes in Figure S1a were from (1) Hi-C data of 30 hours-activated B cell (4DNFIOJNOH8U) (2) HiChIP data of DP thymocytes (GSE141847) and (3) Micro-C of human foreskin fibroblast (4DNFIQXJQWD8). Each example was visualized using Juicebox^18,48^.

### Normalization of chromatin conformation measurement data

Square root vanilla coverage methods^17^ was applied to normalize Hi-C of 30 hours-activated B cell data and HiChIP data of DP thymocytes of C57BL/6J and NOD strains.

### Compartment analysis

Principal component (PC) analysis was performed using HOMER^20^ with 50kb resolution. The regions with positive and negative PC values were regarded as A and B compartment, respectively.

### Protein binding intensity heatmap

The protein binding intensity heatmaps in Figure 2b were generated by our customized R code. The heatmap represents the protein binding intensity on a series of rescaled stripes. Here, each stripe domain is rescaled to 100 bins, and the binding intensities on stripe domain are linearly interpolated. The padding bin sizes are 50 for 5’- and 3’-end of stripe, respectively. The graph on the top of heatmap represents the median binding intensity of each of 200 bins.

### Gene set analysis

The gene ontology analysis of the most highly expressed genes in non-stripy and stripy TAD of mouse DP thymocytes were performed using MetaScape^29^. Both input and analysis species are set as Mus musculus.

### Stripiness comparison between two mouse strains

The stripes from NOD and C57BL/6J DP thymocyte HiChIP data were compared. First, the stripes were searched from both 5kb and 10kb resolution data, and then filtered based on median *P*-value <0.1. Stripes from four datasets (NOD/5kb, NOD/10kb, C57BL/6J/5kb and C57BL6J/10kb) were then merged as follows. First, stripes from same strain but different resolution data were combined. For overlapping stripes, stripes with largest stripiness were selected. Next, stripes from different strains were combined. For combined stripes, stripiness and median *P*-values were recalculated based on NOD and C57BL/6J HiChIP data using Stripenn’s *score* function.

### Statistical analysis

Statistical significance was tested using two-sided Wilcoxon rank sum test. * *P* < 0.05; ** *P* < 0.01; *** *P* < 0.001; **** *P* < 0.0001.

---

**Figure S1.**
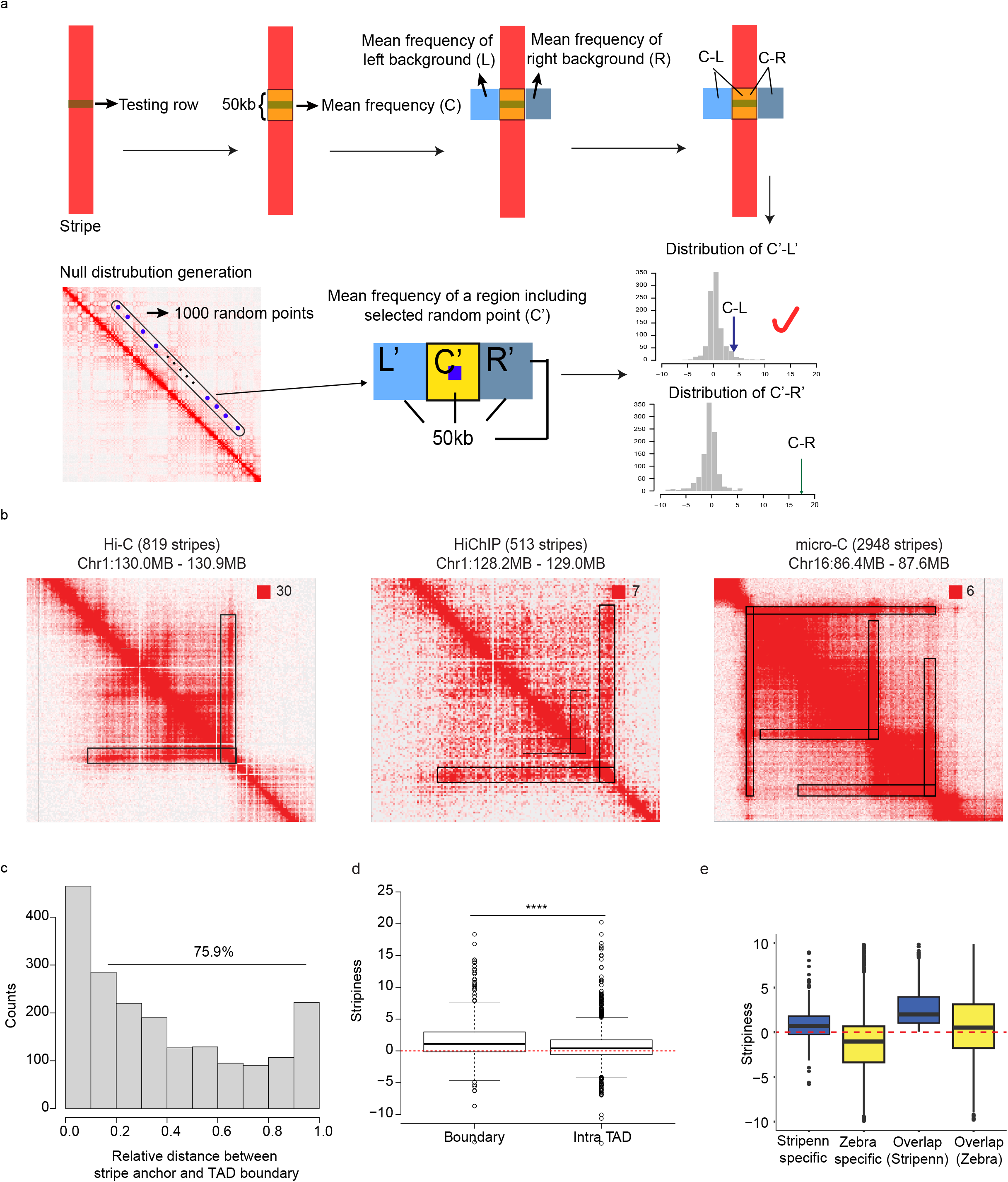
Stripenn detects stripes from various chromatin conformation capture data and has its own scoring system. (a) Description of how *P*-value of individual row in a stripe is estimated. For a testing row (marked green), the mean contact frequency (*C*) is calculated for N rows (yellow) corresponding to 50kb including the testing row at center. The mean frequency of left (L, sky blue) and right (R, blue) adjacent backgrounds are also calculated. Here, the background size is N × N bins. Then, the significance of *C-L* and *C-R* are estimated based on the null distribution of contact frequency difference. To construct null distribution, 1000 random points were selected from which bin distance is identical to that of stripe anchor and testing row. Here, null values for center (C’), left (L’) and right (R’) mean contact frequency is calculated from N x N matrices, respectively. The significance of C-L and C-R are estimated based on null distribution, and less significant value is selected. (b) Example stripe calls (marked black border) from Hi-C (B-cell), HiChIP (T-cell) and Micro-C (human foreskin fibroblast) data. Stripenn detected 819, 513 and 2948 stripes from three data, respectively (median *P*-value<0.05). (c) Histogram demonstrates relative distance between stripe anchors and TAD boundaries obtained from T-cell HiChIP data. For 5’-stripe (3’-stripe), the relative distance is the distance between the stripe anchor and 5’-end (3’-end) of TAD boundary where the anchor is included divided by the TAD size. More than 70% of stripes showed relative distance >0.1. (d) Stripiness comparison between boundary stripe (relative distance < 0.1) and intra TAD stripe (relative distance >= 0.1). (e) Stripiness comparison between Stripenn (blue) and Zebra (yellow). The stripiness distribution of overlapping stripes are different between Stripenn and Zebra because the stripe coordinates are not identical between two methods. Stripiness of overlapping stripes are shown in Figure 1f.

**Figure S2.**
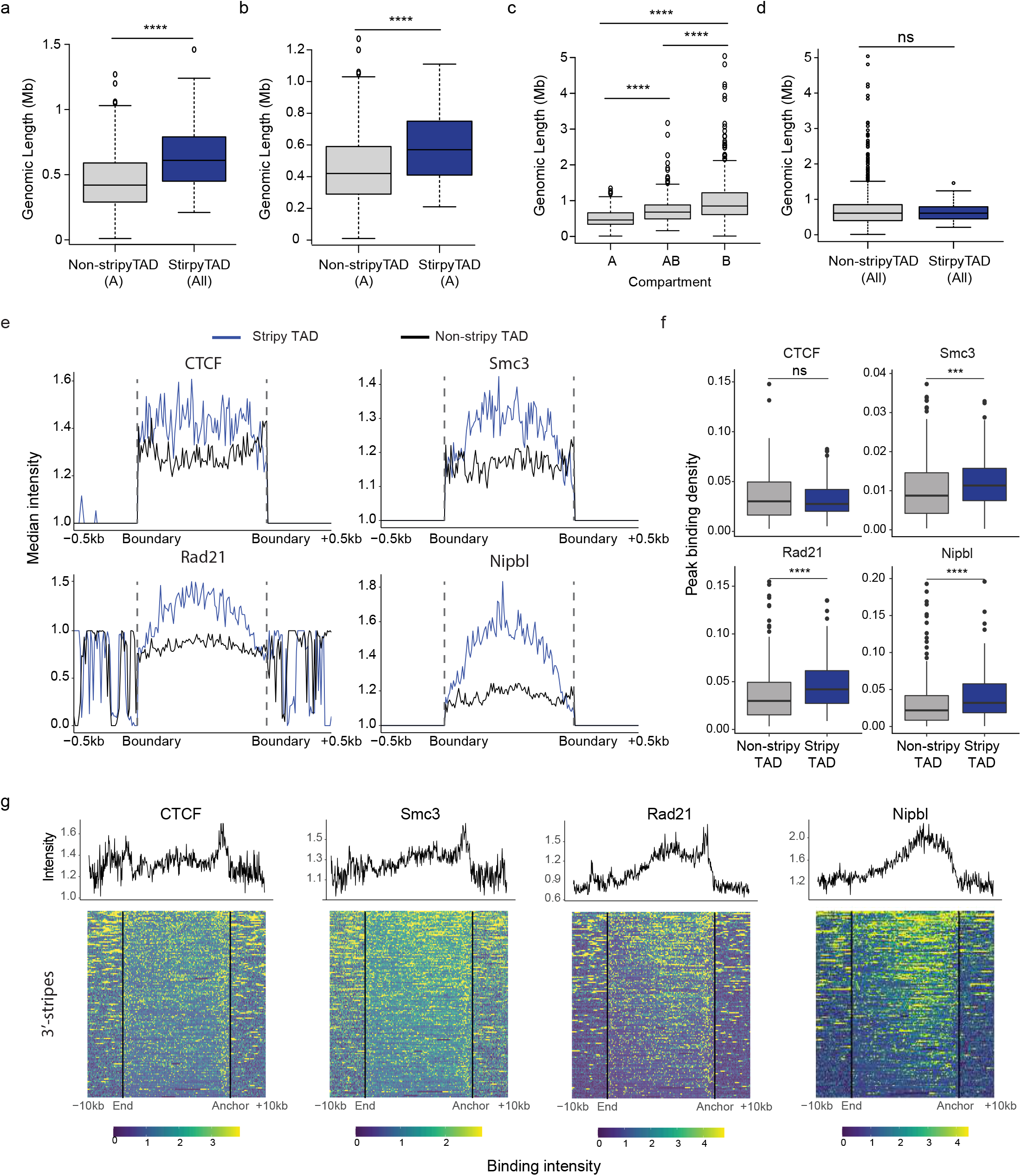
Comparison between stripy and non-stripy TADs using Hi-C data. (a-d) Genomic size distribution of (a) non-stripy TADs in only A compartment and all stripy TADs, (b) non-stripy and stripy TADs, both included in only A compartment, (c) TADs only in A, across A and B (AB) and only in B compartment. (d) all non-stripy TADs and stripy TADs, (e) Median binding intensity of CTCF, Smc3, Rad21 and Nipbl on rescaled stripy TAD (blue) and non-stripy TAD (black). Dashed lines represent the TAD boundary, (f) The structural protein peak binding density in stripy (blue) and non-stripy (gray) TAD. For each TAD, the genomic size of total protein binding sites at peak was divided by the TAD size, (g) Distribution of CTCF, Smc3, Rad21 and Nipbl on rescaled 3’-stripes. The profiles represent the median intensity of each proteins. Black lines in each heatmap are boundary of stripes.

**Figure S3.**
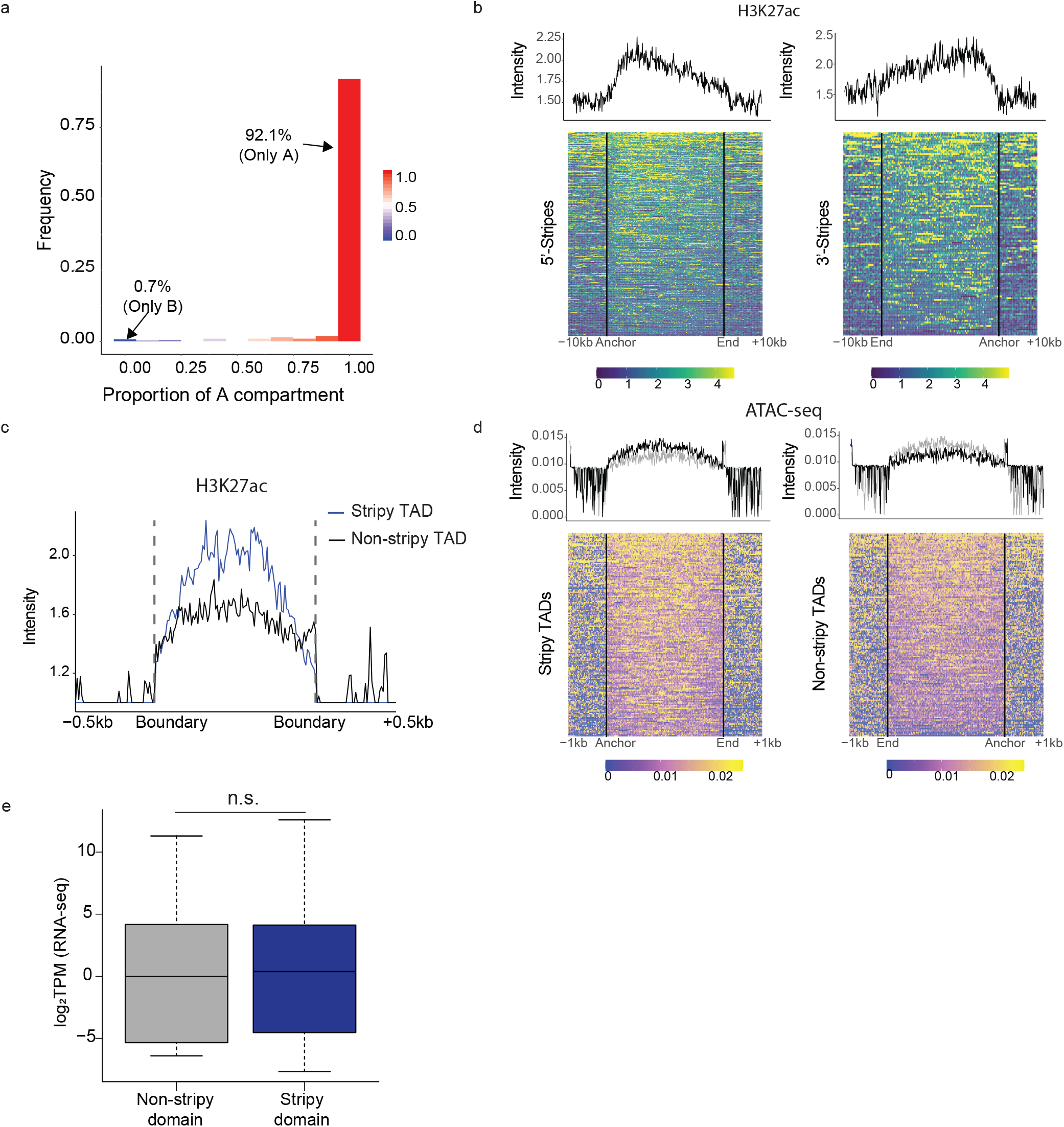
Stripy TADs are epigenetically more active and accessible. (a) Proportion of A compartment within stripe domains from T-cell HiChIP data. 92.1% of stripes are solely in A compartment. (b) Binding intensity of histone mark (H3K27ac) along rescaled stripes from HiChIP data. Left and right plots correspond to 5’ and 3’-stripes, respectively. Yellow pixels represent high intensity. (c) Median binding intensity of H3K27ac on rescaled stripy (blue) and non-stripy (black) TADs. Dashed line represents the TAD boundaries. (d) Comparison of DNA accessibility between stripy TADs (left) and non-stripy TADs (right). The heatmaps represent the degree of DNA accessibility. Solid black lines show TAD boundaries. Black line in upper graph is the median intensity of each column in the heatmap. Gray line is the median intensity of counterpart and added for comparison. (e) Gene expression comparison between non-stripy TAD (gray) and stripy TAD (blue). Here, the expression of all genes within TADs were measured.

**Figure S4.**
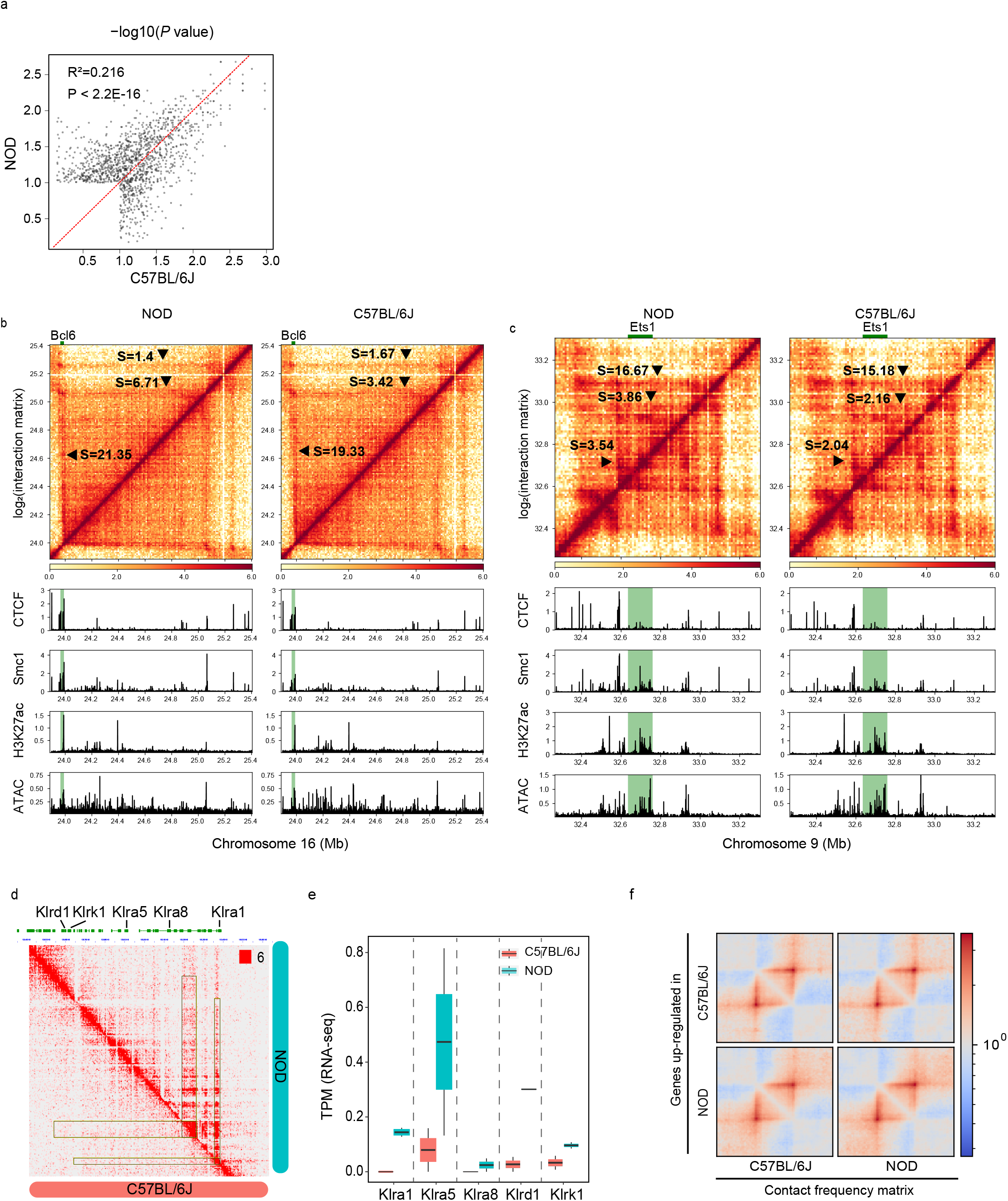
The effect of natural genetic variation on stripe. (a) Stripes from DP thymocytes HiChIP of NOD and C57BL/6J (median *P*-value <0.1) were merged and median *P*-value was recalculated based on HiChIP measurements in each strain. Median *P*-values from two strains show significant correlation (*P*-value < 2.2e-16). (b-c) Stripes and binding intensity of CTCF, Smc1, H3K27ac and Tn5 transposase at (b) *Bcl6* and (c) *Ets1* loci. Stripes are marked a triangles. Gene loci are highlighted with green shadow. (d) Differential stripe between NOD and C57BL/6J at killer cell lectin-like receptor (KLR) gene family loci. (e) Expression level of KLR gene family in C57BL/6J and NOD. (f) Pileup plot of stripes harboring differentially expressed genes.

## Acknowledgement

We thank the Vahedi lab for helpful discussions and support, particularly: Aditi Chandra, Wenliang Wang, and Naomi Goldman. This work was supported by R01HL145754, U01Act DA052715, U01DK127768, the Burroughs Wellcome Fund, the Chan Zuckerberg Initiative, W. W. Smith foundation and the Sloan foundation awards to G.V.

## Notes

### Competing Interest Statement

The authors have declared no competing interest.

